# Three-dimensional single cell patterning for mid- to high-throughput studies of tumor cell and extracellular matrix heterogeneity

**DOI:** 10.1101/163881

**Authors:** Xiangyu Gong, Kristen L. Mills

## Abstract

Cancer heterogeneity includes cancer cell genetic heterogeneity and heterogeneity in the tumor microenvironment (TME)—both stromal cells and extracellular matrix (ECM). Determining which combinations of the vast array of possibly interacting heterogeneities drive tumor progression presents a major multi-disciplinary challenge in cancer research. To make effective treatment decisions this challenge must be addressed. The practical challenge is assessing cellular heterogeneity in a statistically powerful way with respect to targeted TME characteristics. Here we present a simple, but extensible, and low-cost method for conducting mid- to high-throughput and long-term studies of heterogeneous cell responses to various biomechanical stimuli in 3D models mimicking the biomechanical properties of the ECM. Using a platform we term “the drop-patterning chip” thousands of cells are simultaneously transferred from microwells and fully embedded, only using the force of gravity, in precise patterns in 3D. This method allows for throughputs approaching flow-through methods, which lack phenotypic information on cell-matrix interactions, and does not rely on expensive or harsh patterning forces, which often times also result in a proximal stiff surface.

## 1. Introduction

Heterogeneity is a major challenge facing cancer researchers, and a multi-disciplinary discussion is currently underway on how to best address it.^1^ Multiple technologies exist that address the challenge of measuring the heterogeneity of a large population of cells in a statistically powerful way. These include, for example, fluorescence activated cell sorting and microfluidics, both capable of rapid, high-throughput analyses of cells in a flow-through manner. For example, a plethora of cell surface proteins can be recognized and measured that may reveal glioblastoma heterogeneity,^2^ or the optical deformability of a cell can be measured that may reveal its biophysical state,^3^ respectively. Although powerful, these methods are not ideal for assessing cell interactions with the tumor microenvironment (TME), as it is not concurrently possible test cells adhered within a 3D matrix.

When cells are within a 3D matrix, it has been shown that their morphology and differentiation are considerably more representative of *in vivo* conditions as compared to 2D tissue culture (Figure S1).^4^ The reason for this is the cell response to biomechanical stimuli imposed by the ECM. One way in which heterogeneities of cancer cells and the tumor microenvironment may interact is the way in which individual cancer cells respond to altered biomechanical stimuli. Biomechanical stimuli, controlled by the tissue-specific composition of the ECM and its disease state, include mechanical forces and matrix stiffness. It is becoming increasingly clear that experiments intended to accurately assess phenotypic cell response to biomechanical stimuli should be conducted in 3D matrices that recapitulate the morphology of the native tissue as well as its mechanical properties.^5–7^ Current methods to measure cell response to biomechanical stimuli in 3D are unable to quantify the heterogeneity of a large population of cells. They are either based on direct observation of small numbers of cells,^5^ which does not allow for statistical power, or based on large population assays,^8,9^ such as Western blots, which mask heterogeneity.

Towards gathering large sets of data on single, attached cells, investigators have devised patterning methods and scaffolds that array single cells or small numbers of cells on various substrates and surface morphologies. Cell placement has been achieved via microwells^10–13^ or surface patterns in which small areas of the substrate are modulated to promote cell attachment and elicit a mechanobiological response.^14–17^ Microwells have not only been used as a niche where cells may proliferate, but they have also been used as a tool for transferring cells into another 2D environments.^18,19^ Others have used engineered scaffolds, such as polymer structures fabricated via direct laser writing (DLW)^20^ and crack-based patterning^21^ to provide single cells adhesive, topological supports in a 3D space. Whereas these methods allow for cell anchorage and ease of image collection and analysis, the stiff and/or 2D nature of the substrates do not provide an accurate analog to the soft, 3D nature of the *in vivo* environment.

Patterning cells that are completely embedded within a 3D, uniform, and soft environment while enabling high-throughput studies presents a particular challenge. Some innovative methods towards achieving this include dielectrophoretic (DEP)^22–24^ or magnetic^25^ forces or surface acoustic waves (SAWs)^26,27^ to manipulate cells into patterns. Position control over cell placement is indeed accurate, however these methods require special tools (e.g., gold coated nodes, SAW chips) not easily accessible in every lab. Furthermore, although viability is confirmed after patterning, the forces being applied could potentially be harmful to the cells or alter their expression.^28^ The major drawback of these methods for biomechanical experiments is the presence of stiff substrates necessitated by the patterning methods, which prevents the full encapsulation of the cells or cell clusters in an ECM-mimicking matrix.

Fully embedding cells in a 3D matrix is most simply achieved by mixing cells with a liquid precursor to a synthetic or biological hydrogel and allowing the gelation process to encapsulate the cells. Long-term monitoring of selected single cells or cell clusters in a mid- to high-throughput fashion then becomes a significant challenge, if not impossible, as the cells are positioned randomly. For long-term studies of single-cell behavior, researchers have resorted to embedding small numbers of cells into a matrix,^5^ which eases the experimentalist’s efforts to locate cells, but often does not provide a large enough sample set for significant statistical analyses.

No method has yet been devised in which single cells or small cell clusters may be accurately patterned in a 3D environment allowing for high-throughput studies of heterogeneous cell responses to biomechanical stimuli. The goal of this work was to develop a simple and low-cost platform for patterning a large population of single cells or cell clusters completely within a 3D environment (without any undesirable discontinuous or stiff boundaries). The resulting platform was to allow for longitudinal observation of the single cells or cell clusters and control over the cell-cell distance within various cell array patterns. In this article, we describe the design, fabrication, method, and performance of the platform that we developed.

## 2. Results

Our platform capitalizes on the single-cell arraying power of microwells fabricated by the standard soft lithography technique. Once the cells are trapped in the microwells, it is inverted and the cells are allowed to drop into a hydrogel that is in the process of gelation. Therefore, we term the platform the drop-patterning chip (Figure 2a). The detailed operation of the drop-patterning chip is first presented for large-scale arrays. Performance of the method is evaluated by its patterning efficiency, single-cell occupancy rate, and spatial distribution of cells. The gentle transfer method furthermore leads to excellent cell viability. Finally, we present longitudinal studies of cell motility and *in situ* tumor development from these individual cells.

### 2.1. The drop-patterning chip

The drop-patterning chip consists of three layers (Figure 1a) creating a closed chamber, which may be filled with hydrogel (Figure 1b). The three layers, from bottom to top, are a PDMS microwell substrate, a square frame PDMS spacer, and a standard glass coverslip. Due to the natural adhesion of PDMS to itself and between PDMS and glass, the three layers spontaneously bond together with slight pressure applied by hand. The microwell dimensions—30 μm in diameter and 27 μm in depth—were determined by experimentation to most efficiently capture one cell per well. All data presented here are from experiments carried out on microwell arrays with center-to-center spacing (along both columns and rows) of 100 μm, 150 μm, or 200 μm.

**Figure 1.**
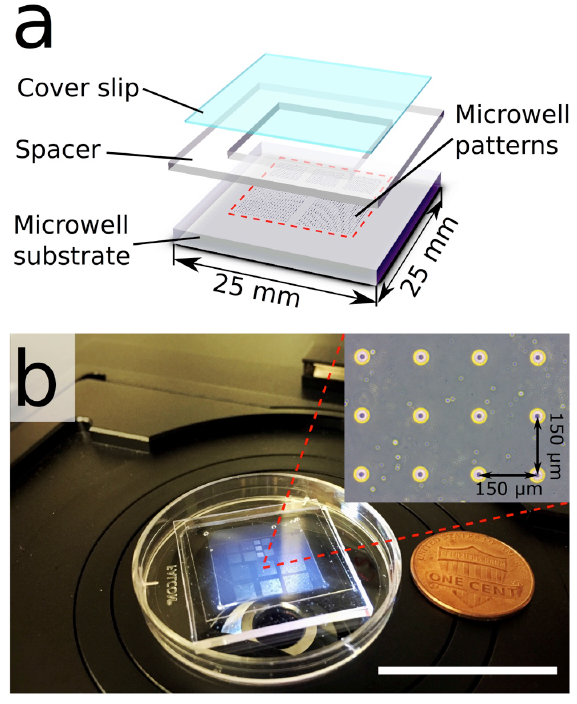
Drop-patterning chip. (a) Schematic of the three-layer configuration of the drop-patterning chip. From bottom to top: PDMS microwell substrate (1. 5 mm thick), square frame PDMS spacer (inner side lengths 15 mm, thickness: approximately 600 μm), standard glass cover slip (22-mm square) (b) Photograph of an assembled chip. The chip was designed to fit in a 35-mm Petri dish lid. The inset shows part of the microwell arrays with inter-well spacing (along both columns and rows) of 150 μm. (Scale bar: 25 mm)

**Figure 2.**
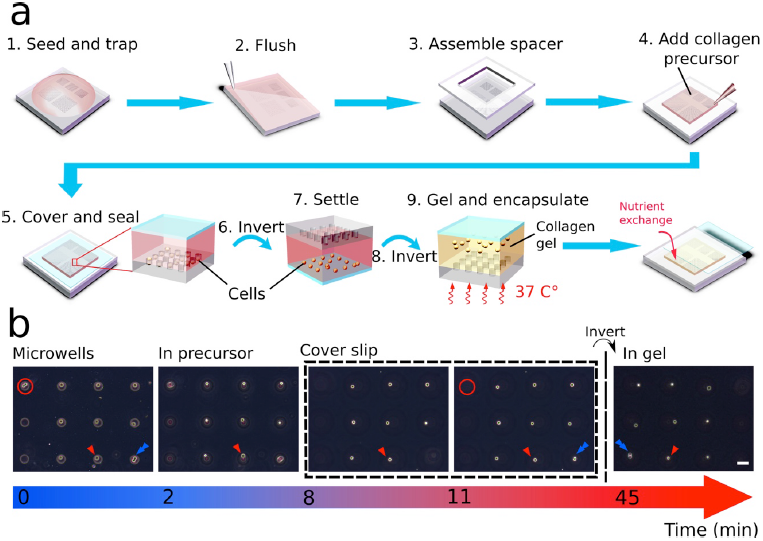
Schematics and images to illustrate the drop-patterning method. (a) 1. A cell suspension is seeded on a substrate containing arrays of microwells. The cells are allowed to settle for 5 minutes. 2. Excess cells are gently flushed away with PBS. 3. A PDMS spacer is assembled onto the microwell substrate to create a chamber. 4-5. Once filled with collagen solution, the entire chamber is sealed with a cover slip. 6. The chip is inverted and the trapped cells begin falling out. 7-9. When most cells settle on the glass cover slip, the chip is inverted again and the collagen is allowed to gel at 37 C°. After the cell pattern is embedded in collagen gel, the cover slip is carefully slid aside for nutrient exchange. (b) On chip, real-time monitoring (bright field imaging) of cell array embedding in collagen by drop-patterning. (Movie S1) The imaging started immediately after the first inversion, and always focused on a single cell indicated with a red triangle. The cell doublet in the red circle was stuck in the well and did not fall into the collagen precursor. The blue arrow may be used to keep track of a cell/position before and after the second inversion. Enlarged and enhanced images of the cells before and after collagen gelation are presented in Figure S3. The inter-well spacing is 150 μm. (Scale bar: 50μm)

### 2.2. Step-by-step drop-patterning in collagen

The first set of steps in the drop-patterning technique includes priming the microwells with cells and assembling the chip (Figure 2a, steps 1-5). A cell suspension was deposited on to the PDMS substrate patterned with microwell arrays. Treating the PDMS substrate with plasma and then bovine serum albumin (BSA) prior to seeding promoted wetting and prevented cell attachment, respectively. After allowing the cells to settle into the microwells for 5 minutes, excess medium and cells were removed from the surface with a gentle phosphate-buffered saline (PBS) wash. The PDMS spacer (middle component, Figure 1a) was then assembled atop the substrate, creating a chamber. This chamber was filled with a collagen I solution (1.0 mg/mL) and then sealed with the coverslip (top component, Figure 1a)

The second set of steps in the drop-patterning technique transfer the array of cells into the collagen gel (Figure 2a, steps 6-9). The enclosed chip was inverted and the trapped cells were allowed to fall, due to gravity, out of the microwells. Adhesion of the cells to the PDMS microwell walls varied, therefore not all cells fell out of the microwells simultaneously. This means pattern fidelity may be lost and some cells may not be fully transferred into the gel. Therefore, we first let the cells fall all the way to the coverslip. The chip was warmed at 37 °C in an incubator for 4 minutes and 30 seconds, allowing the collagen to start gelling. Then we inverted the chip once again upon which time all cells fell from the coverslip simultaneously, due to the relative uniformity of adhesive interaction between the cells and the BSA-treated coverslip. The synergy of the rate of collagen gelation and velocity of the falling cells resulted in the cell array becoming fully encapsulated.

To visualize the drop-patterning method, the whole process was monitored under a bright field microscope (Zeiss, Axio Vert.A1) (Figure 2b and Movie S1). Focus was maintained on a single cell (indicated with red triangles) in the array. Immediately after the first chip inversion (labeled Time 0), the cells were still trapped in the microwells. After 2 minutes, the microwells were out of focus, which means cells left the microwells and were traveling through the collagen solution. It took approximately 8 minutes for all single cells and 11 minutes for a doublet of cells (indicated with a blue arrow) to settle on the glass coverslip. Another doublet remained stuck in a microwell (indicated with a red circle). After an incubation time of 4 minutes and 30 seconds at 37 °C, the chip was inverted a second time and incubated at 37 °C for 30 minutes longer. The resulting fully embedded 3 × 3 cell array is shown in the final image of Figure 2b (Figure S2).

### 2.3. Cell patterning efficiency

Trapping efficiency is considered an important measure of the performance of microwell arrays in high-throughput research on anchorage dependent cells.^11,29^ Similarly, the percentage of cell-occupied positions in the embedded 3D pattern reflects the overall efficiency of the drop-patterning method, which we defined as cell-patterning efficiency. As shown previously, not all trapped cells fell out of the wells, which means there was a nominal discrepancy between the cell-trapping efficiency in the microwells and the cell-patterning efficiency in collagen. Both the cell-trapping efficiency and cell-patterning efficiency were calculated according to equation 1, by dividing the total number of occupied positions either in the microwell pattern or in the collagen by the total number of microwells.

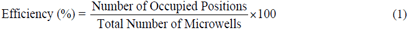

Cell-trapping and cell-patterning efficiencies for this method were measured on a total of eighteen 20 × 20 microwell arrays (e.g., Figure 4a) on two chips. Three different inter-well spacings were used: 100 μm (8 arrays), 150 μm (5 arrays), and 200 μm (5 arrays) (Figure 3a). No statistical difference occurred in the efficiencies between the different inter-well spacings according to two-tailed student t-tests (p > 0.05). The average cell-trapping efficiency and cell-patterning efficiency of all the arrays were 75% and 62.3%, respectively.

**Figure 3.**
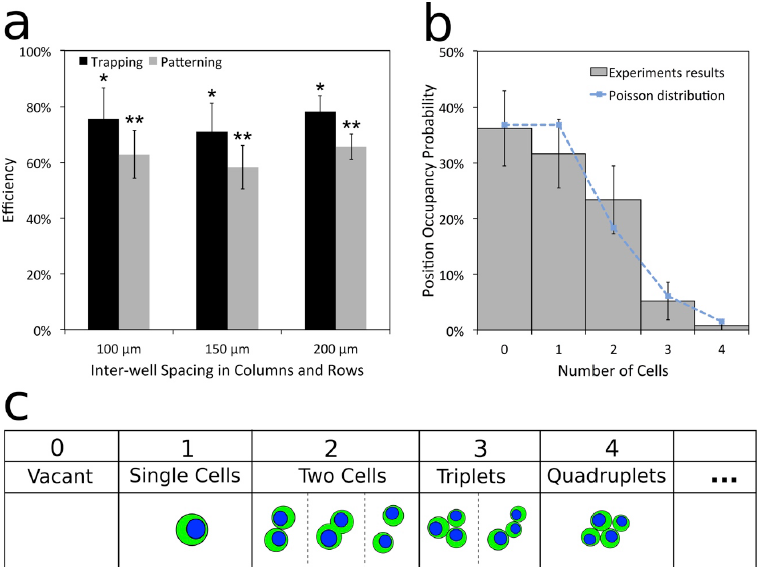
(a) Trapping efficiency and patterning efficiency of arrays with inter-well spacing of 100 μm (n = 8 arrays), 150 μm (n = 5 arrays), 200 μm (n = 5 arrays). No statistical difference occurred in the efficiencies between the different inter-well spacings according to two-tailed student t-tests (p > 0.05). (b) The probability distribution (histogram) of number of cells per position determined from observations at 6,000 positions of fifteen 20 × 20 arrays (n = 15). The theoretical Poisson distribution is overlaid with a dotted line. (c) The schematics illustrate the observed cell occupancy scenarios at the drop-patterned positions.

**Figure 4.**
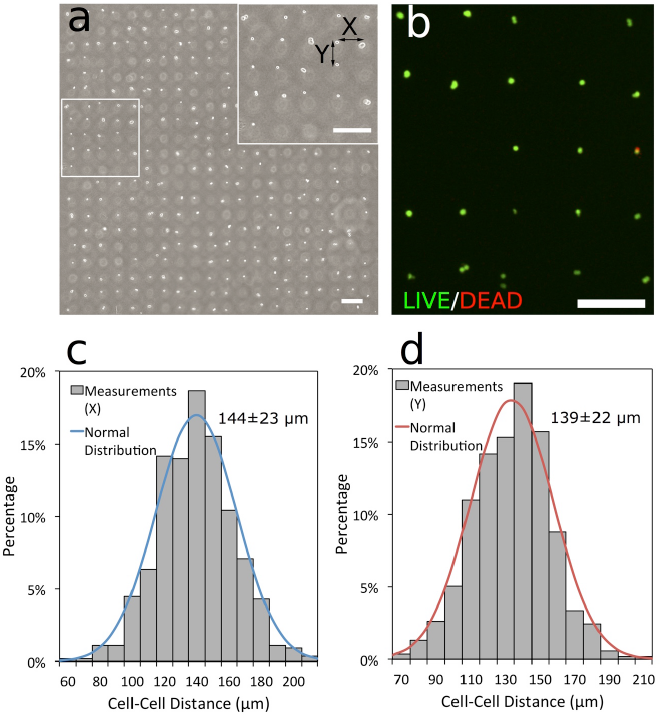
Large-scale cell array patterning in single-cell resolution. (a) Cell array patterned in collagen by 20 × 20 microwell array with ll spacing of 150 μm in the columns and rows. The embedding efficiency is 65. 8%. (b) Viability assay (19 hours after embedding) of drop patterning. A cell array (5 × 5 positions, spacing: 200 μm, efficiency: 88%) was stained with calcein AM (live cells, green) and EthD-1 (dead cells, red). (c) Distribution of cell-cell distances in the rows (X) as a measurement of drop patterning precision. (d) Distribution of cell-cell distances in the columns (Y). 536 measurements were done for each direction, over three arrays (nominal spacing: 150 μm) on two chips. (Scale bars: 200 μm)

In addition to single cells, small cell clusters, such as doublets and triplets, were trapped in the microwells and patterned in collagen (Figure 3c). This occurred due to the variation of cell size and cell-cell adhesion. The probability of finding a given number of cells in a position corresponding to the 20 × 20 positions of an array (n = 15 arrays) is presented in Figure 3b, where it may be seen that the single-cell occupancy rate was 31.6%. The resulting distribution is dictated by Poisson statistics (with λ = 1), which has been shown with other cell trapping methods.^29,30^

## 2.4. Characterization of spatial distribution

While cells are being drop-patterned in the collagen solution, drifting in the plane of the array is inevitable. Drifting influences the patterning fidelity (Figure 4a). To quantify the fidelity of the drop-patterning method, we measured the horizontal (X) and vertical (Y) distances (arrows in Figure 4a) between two cells or clusters that were patterned by two neighbor microwells with the spacing of 150 μm. Figures 4c and 4d are the histograms showing the cell-cell distance distribution over 536 measurements in each direction. The mean cell-cell distance was 144 ± 23 μm in X and 139 ± 22 μm in Y. This suggests that the resolution of the drop-patterning method is approximately 20 μm.

### 2.5. Patterned cell viability

In order to confirm that drop-patterning does not damage living cells, a cell viability assay was conducted via calcein acetoxymethyl ester (AM) and ethidium homodimer (EthD-1) staining 19 hours after the cells had been patterned in the collagen gel (Figure 4b). In four random regions of a drop-patterning chip, 361 cells were imaged. Only three cells had died (were fluorescing red), which indicates cell viability may be 99% when following the drop-patterning protocol.

### 2.6. Longitudinal study of cell motility, division, and model tumor development

The drop-patterned arrays enable the study of the heterogeneity of behaviors in 3D of single tumor cells over longer periods of time. An array of colon cancer cells (HCT-116) was drop-patterned into collagen and their progression was tracked in nine positions over 3.5 days (Figure 5a). Here, we demonstrated the ease of following the progression of multiple cells over periods of time when a sample is not kept directly on a microscope stage equipped with an incubation system. Because the fully gelled collagen detached from the PDMS microwell substrate after we added cell culture medium, we noticed a position shift between the embedded cells and their corresponding microwells underneath. But that did not interfere with locating these cells. For illustration, the 3 × 3 cell array has been visually divided and labeled as zones I through IX. The patterned array consisted of eight single cells (I-VIII) and one doublet (IX). All cells survived and divided within two days.

**Figure 5.**
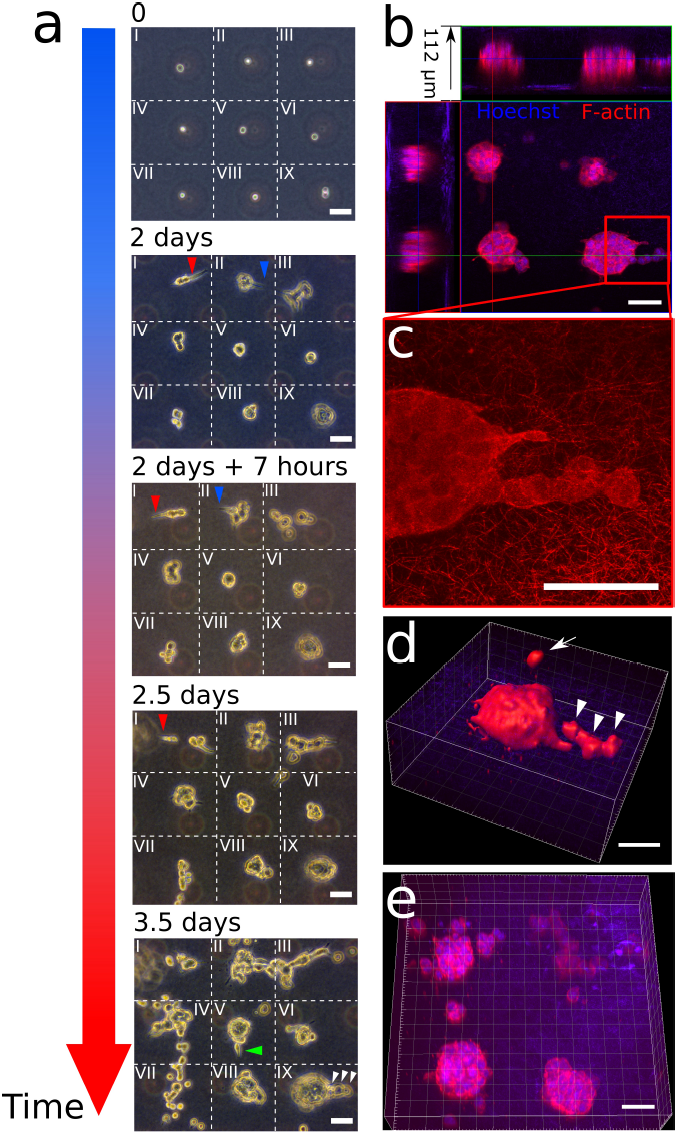
3D on-chip cell motility and tumor development longitudinal observations. (a) Phase-contrast images of tumor progression from a 3 × 3 cell (HCT-116) array in collagen (1.0 mg/mL). The field of view is divided into nine zones (I-IX) with dotted lines. Cell protrusions over time are indicated with arrowheads (red: zone I, blue: zone II, green: zone V, white: IX). (Scale bars:50 μm) (b) Immunofluorescence imaging (red: F-actin, blue: Hoechst) of a tumor array developed from the cells in (a) (Scale bar: 50 μm). (c) Confocal reflectance imaging shows the collagen I (1.0 mg/mL) microstructure (thickness: 7 μm, maximum projection), and verifies the tumors were fully embedding when developing (Scale bar: 50 μm). (d) 3D rendering of the bottom right tumor in (b) showing single cell migration (arrow) and collective migration (arrowheads) (Scale bar: 40 μm). (e) 3D reconstruction of a tumor array visualizing tumor shape and cell organization (Scale bar: 40 μm).

Cell protrusions (Figure 5a, red and blue arrowheads) and motility were readily followed over the observation time period. Notably, the cells in zones I and II exhibited extensive protrusive behaviors into the collagen matrix starting from day 2, with the protrusions changing directions during the first 7 hours of day 2. Also in the first half of day 2, cells from zones I, III and VII underwent phenotypic change resembling epithelial-mesenchymal transition (EMT) and migrated away from their respective initial positions. After 3.5 days in the collagen gel, cells in zones II, III, IV and VII displayed the highest motility. They continued to proliferate and migrate into neighboring zones.

In contrast, the cells in zones V, VI, VIII and IX were not, at least initially, as motile, however, they did proliferate. The three single cells and one cell doublet began forming multicellular tumor spheroids within the first two days. Only after three days did cells from these tumor spheroids begin to form protrusive actions and migrate away from the tumor spheroids, apparently in both single-cell (green arrowhead) and collective (white arrowheads) manner.

After 3.5 days, the cells in the 3 × 3 array were fixed and their nuclei (Hoechst, blue) and F-actin (phalloidin, red) were stained. A confocal image stack (height: 112 μm) was acquired from a focus plane above a tumor array to a focus plane below the tumor array (Figure 5b). Collagen fibers were auto-fluorescent in the blue channel on both the first and the last several images with no F-actin (red) observed, which validated that all the tumors were fully embedded in collagen. Reflectance confocal microscopy was used to visualize detailed microstructure of collagen in 7-μm thickness (Figure 5c). 3D-reconstruction of a solid tumor (Figure 5d) and a tumor array (Figure 5e) clearly shows the spatial relationship between tumors and their neighboring cells. Figure 5d clearly shows that cell migration did not only happen on the focal plane. A single cell escaped the tumor and migrated upwards (marked with an arrow).

### 2.7. Other single cell resolution patterns in 3D

Array patterns may be an efficient tool for high-throughput quantitative studies of the heterogeneity of single cell behaviors in 3D. However, the drop-patterning method may be used to produce other, more complex patterns that may be designed, for example, to induce desired cell-cell interactions by spatially variations in the cell density. We demonstrated this capability with concentric-circle patterns and the representation of the letters R, P, and I (abbreviation for Rensselaer Polytechnic Institute) in Figure 6. In the concentric-circle pattern, the cell density varies along the radius and the cell-cell distance can be varied within one pattern (arrows). The inner region, with denser cell population, resulted in more cell-cell interaction and two tumors were observed to merge into one (red circles). Figure 6c and d show fluorescence images of the concentric-circle patterns in Figure 6b and capital letters R, P, and I developed *in situ* for 6 days from single cells or small cell clusters drop-patterned into collagen.

**Figure 6.**
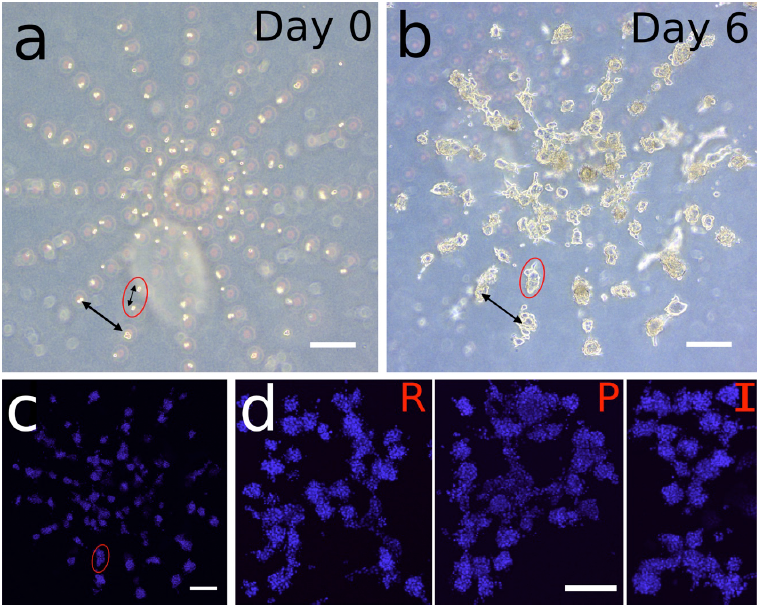
Various patterns of single cells in collagen with the drop-patterning method. (a) Single cells and small cell clusters were drop-patterned in collagen in concentric circles demonstrating the capability of producing varying intercellular spacing within one pattern (arrows). Ninety-nine of total 128 positions (78% patterning efficiency) were occupied by cells. (b) Six-day culture of the cell pattern in (a) showing tumor growth, cell migration, and cell-cell interactions. Red circles indicate two separate cells growing and merging into one tumor after six days. (c) A fluorescence image (Nuclei: Hoechst, maximum projection) of the cell pattern in (b). (d) Fluorescence images (Hoechst, maximum projection) of cell-assembled capital letters R, P, and I (abbreviation for Rensselaer Polytechnic Institute) after six-day growth in the collagen. (Scale bars: 200 μm)

## 3. Discussion and Outlook

Among other measurements, we have evaluated the drop[[[[-patterning method by its trapping, patterning, and single-cell efficiencies. The state of the art with similar microwell trapping technologies produces trapping efficiencies as high as 85%-92%.^10^ Our experience, however, and that of other studies indicate that the single-cell trapping efficiency of passive methods such as this is highly dependent on cell type and the ease with which the cells are separable in suspension. This results in single-cell efficiencies on the order of 26%^31^ to 34%,^29^ and probabilities of cell numbers per well or per chamber fitting the Poisson distribution.^30^ Our single-cell efficiency is currently on par with these other studies, however, this may significantly be improved by incorporating the drop-patterning method into a microfluidic platform to trap the cells more efficiently.^32,33^

Technologies, such as cell encapsulation^34^ and 3D gel-islands in a microfluidic channel^29^, are able to provide cells a 3D environment while allowing for cell proliferation to be tracked in individual droplets or chambers. However, the ability to expand the complexity of the micro-scale geometrical, chemical, or biological composition of the matrix is limited and cell-cell interactions are precluded. The drop-patterning technique allows for a large degree of flexibility in experimental design, both in terms of the heterogeneity of interactions between cells and the ECM and in the pattern geometry. The results presented here have focused on collagen I hydrogel as the matrix, but any hydrogel formulation for which the gelation kinetics may be controlled on the time scale of minutes may be used. This provides the basis for which cell-matrix interactions may be studied with respect to many different biochemical compositions, morphologies, and stiffness of ECM in a statistically powerful way. Furthermore, a large variety of cell-cell interactions would be made possible through the design of the inter-well spacing. These include mechanical or diffusive cell-cell communication.

Heterogeneity of cancer cell interactions with the TME extends beyond cancer cell-ECM interactions to cancer cell-stromal cell interactions. The drop-patterning platform additionally paves the way to enable ease of creating high-throughput 3D co-culture studies with, for example, cancer associate fibroblasts, endothelial cells, or immune cells.

Although single-cell microwells and arrays were used here to illustrate the use of this new tool, we would like to point out that it is possible to tune the size of the wells in order to trap aggregates of cells or tumor spheroids.

The flexibility with which one may choose the cell patterns as well as the types of hydrogel lays the foundation for tissue engineering, stem cell research, *ex vivo* cancer assay, and many *in vitro* studies on intercellular communication (e.g., neurite outgrowth of primary neurons) and tumor-microenvironment interaction (e.g., tumor angiogenesis). The drop-patterning method may be easily extended into a multiplex, co-culture platform, integrating with other platforms, such as a chemotaxis device^35^ and a engineered blood vessel^36^, to create a complex engineered niche for high-throughput studies of tumor progression.

## 4. Materials and methods

### 4.1. Design and fabrication of the drop-patterning chip

We designed the microwell patterns on computer aid design (CAD) software SolidWorks (Dassault Systèmes), and all the patterns were printed on a chrome mask by a high-resolution printing service (Front Range Photo Mask, CO, USA). Micro-posts on the silicon wafer were fabricated in a negative photoresist (SU-8 3025, MicroChem, MA, USA) through the techniques of photolithography. In brief, a layer of SU-8 (27.5 μm thick) was spin-coated on a 3-inch silicon wafer for 30 sec at 2,800 rpm. After soft baking at 95 °C for 10 min, the SU-8 coating was crosslinked under the exposure of UV light through a chrome mask. After post-exposure baking at 65 °C for 1 min and 95 °C for 8 min, non-crosslinked SU-8 photoresist was washed off through developing process, and all the crosslinked micro-posts remained on the silicon wafer. The silicon wafer was then hard baked at 180 °C for 10 min. The heights of these SU-8 features were inspected by a stylus profilometer (Veeco, DekTak 8). Using soft lithography techniques, we molded microwells onto a PDMS (Sylgard 184, Dow Corning) sheet via the silicon master with micro-posts. First, we treated the surface with Tridecafluoro-1,1,2,2-tetrahydrooctyl-1-trichlorosilane (TFOCS, Gelest, PA, USA) to prevent cured PDMS sticking to the master. Then, we poured thoroughly mixed, degassed PDMS precursor (ratio of base and curing agent = 10:1 by weight) onto the silicon master in a plastic Petri dish and allowed the PDMS to cure at 70 °C overnight. After the PDMS was fully cured, we peeled it off and cut it into the desired size (25 mm × 25 mm). A PDMS spacer ring (thickness: approximately 600 μm) was directly cut out of a plain PDMS sheet as a square, matching the size of the patterned PDMS substrate. A square coverslip with standard dimensions (22 mm × 22 mm, thickness: 120 – 160 μm) was used to seal the whole device. Before assembling, we treated the PDMS substrate and spacer ring in a plasma cleaner (Harrick Plasma) and allowed them to partially recover its natural hydrophobicity in a sterile ambient environment overnight. All three components were sterilized all the three components with 70% ethanol and then with UV light for 15 minutes.

### 4.2. Preparation of collagen gel

Eight parts of type I bovine collagen monomer solution (3.1 mg/mL, pH 2, PureCol, Advanced Matrix, USA) was diluted with one part of 10× PBS, and then neutralized to a pH of 7.2 – 7.6 with 0.1M sodium hydroxide (NaOH) solution. To avoid local pH variance, the solution was pipetted up and down each time a fraction of NaOH was added. Final volume was adjusted to ten parts with ultrapure water. By this method, the concentration of the neutralized collagen solution was 2.48 mg/mL. Based on the final concentration desired, we were able to adjust the collagen solution to any concentration lower than 2.48 mg/mL, by diluting with cell culture medium. To prevent local gelation, all the solutions and tubes were chilled and all mixing operations were conducted on ice. Since air could be introduced via mixing process, the final collagen solution was degassed on ice in a vacuum desiccator, so that no bubbles formed during gelation.

### 4.3. Cell culture

In the study of cell proliferation and tumor growth in 3D, we chose a human colon cancer cell line HCT-116 (ATCC) as a cell model. Before loading the cells in the chip, we cultured them on tissue culture flasks in McCoy’s 5A modified medium (Corning) with 10% (vol/vol) FBS (Gibco) and 1% penicillin/streptomycin (Gibco) at 37 °C and 5% CO2 in a humidified incubator. The cell culture medium was changed every other day and passaged when cells reached over 80% confluency. When the cells were imbedded in 3D collagen, we continued culturing them by submerging the chip in fresh cell culture medium. Passage numbers of the cells used in this research did not exceed ten.

### 4.4. Device preparation and assembly

Before drop-patterning, the surface of the inherently hydrophobic PDMS microwell substrate (bottom component, Figure 1a) was treated with air plasma (Harrick Plasma) for 30 seconds, a process that renders its surface hydrophilic. The substrate was then placed in a sterile ambient environment overnight to partially recover its natural hydrophobicity. This step allowed for optimal wetting behavior while preventing cell attachment to the microwell walls. The surface of the plasma-treated microwell substrate and the glass coverslip were then incubated at room temperature for one hour with a 10% bovine serum albumin (BSA) solution to further prevent cell attachment.

After the BSA treatment, a cell suspension was seeded on the microwell substrate and the cells were allowed to settle into the microwells for about 5 min. Then the supernatant was removed and excess, untrapped cells were gently flushed away with phosphate-buffered saline (PBS). A Kimwipe was used to carefully dry the unpatterned area near the edges of the substrate, while keeping the central, patterned area wet. A dry PDMS spacer (middle component, Figure 1a) was then rapidly placed onto the substrate to create a chamber. This chamber was filled with a collagen solution (1.0 mg/mL) and then sealed with the coverslip (top component, Figure 1a).

### 4.5. Immunohistochemistry

HCT-116 tumor arrays were grown in 3D collagen for multiple days. The samples were then washed in PBS, fixed with 3.7% paraformaldehyde at 37 °C for 30 min, and permeabilized with 0.5% Triton X-100 at 37 °C for 30 min. After washing with PBS three times for 30 min, the samples were blocked for 10 hours in 5% BSA in PBS at room temperature. An F-actin probe rhodamine phalloidin (1:50, R415, Thermo Fisher) was then applied in dark at 4 °C overnight. After, nuclei were stained with Hoechst (0.2 μg/mL, Hoechst 33342, Thermo Fisher) at room temperature in dark for 4 hours. In the cell viability essay, HCT-116 single cells were stained with calcein AM and EthD-1 (LIVE/DEAD Viability Kit, Invitrogen) after the cells were drop-patterned in collagen for 19 hours.

### 4.6. Image acquisition and statistical analysis

Bright field images of microwells and cells were obtained with an inverted microscope (Zeiss, Axio Vert.A1). Fluorescence images of cells/tumors and reflectance imaged of collagen microstructure were acquired with a laser scanning confocal microscope (Zeiss, LSM 510 META). 3D reconstructions of z-stacks of tumors arrays were performed on software ZEN (Zeiss) and 3D rendering based on confocal microscopic images was made by software Imaris 8 (Bitplane). Cell-cell distances were measured with an image-processing program ImageJ. Statistical analysis was performed on MS Excel. All data was presented as mean ± standard deviation. Statistical difference was determined by two-tailed student t-tests.

## Acknowledgements

The authors acknowledge funding from Rensselaer Polytechnic Institute (RPI) through the School of Engineering, the Department of Mechanical, Aerospace, and Nuclear Engineering. We thank Dr. Aram Chung, Dr. Leo Wan, and Y. Deng for technical assistance. Photolithography for this work was performed in the Micro and Nano Fabrication Clean Room (MNCR) at RPI.

## Author Contributions

X.G. and K.L.M. designed the experiments and wrote the main manuscript. X.G. conducted all the experiments, analyzed data and prepared figures.

